# Coupling optogenetics and light-sheet microscopy, a method to study signal transduction *in vivo*

**DOI:** 10.1101/114686

**Authors:** Prameet Kaur, Timothy E. Saunders, Nicholas S. Tolwinski

**Affiliations:** Division of Science, Yale-NUS College, 12 College Ave West, #01-201, Singapore 138610; Mechanobiology Institute, National University of Singapore 117411,, Singapore; Department of Biological Sciences, National University of Singapore 117543, Singapore; Institute of Molecular and Cell Biology, A*Star, Proteos 138673, Singapore

## Abstract

Optogenetics allows precise, fast and reversible intervention in biological processes. Light-sheet microscopy allows observation of the full course of embryonic development from egg to larva. Bringing the two approaches together allows unparalleled precision into the temporal regulation of signaling pathways and cellular processes *in vivo*. To develop this method, we investigated the regulation of canonical Wnt signaling during anterior-posterior patterning of the *Drosophila* embryonic epidermis. Cryptochrome 2 (CRY2) from *Arabidopsis Thaliana* was fused to mCherry fluorescent protein and *Drosophila* β–catenin to form an easy to visualize optogenetic switch. Blue light illumination caused oligomerization of the fusion protein and inhibited downstream Wnt signaling *in vitro* and *in vivo*. Temporal inactivation of β–catenin confirmed that Wnt signaling is required not only for *Drosophila* pattern formation, but also for maintenance later in development. We anticipate that this method will be easily extendable to other developmental signaling pathways and many other experimental systems.

## Introduction

Dissections of signaling pathways and their downstream processes have centered on *in vitro* cell culture models and *in vivo* model organism studies using small molecule, knockdown and overexpression approaches. Although these approaches have been very useful, they do not lend themselves easily to the study of the spatial and temporal regulation of cell signaling pathways and cellular processes in the dynamic environment of a living organism. Normal genetic crosses using mutations and RNAi techniques are limited in their applications as they result in irreversible knockout or knockdown of a gene, which can be lethal or incomplete ^1^. The recent discovery of optogenetic tools has provided a new and effective toolbox for spatial and temporal regulation of proteins in various *in vitro* and *in vivo* systems ^2-5^. This is made possible by the rapid response of these proteins to light, such as the blue-light mediated activation of *Arabidopsis thaliana* cryptochrome 2 (CRY2) ^6^. We chose to use the CRY2 system as it has proven effective in *Drosophila* embryos ^7^.

Even though the CRY2 system can be activated simply with 488nm lasers, there is high phototoxicity and bleaching which prevent collection of temporal developmental information. Hence, the application of optogenetic tools is only as good as the visualization approach that can be brought to bear. The latest advances in light-sheet microscopy facilitate high resolution optical sectioning and live imaging of embryos with low phototoxicity as compared to conventional laser confocal imaging ^8^. Importantly, light-sheet microscopy enables selective activation of optogenetic insertions without significant out-of-plane effects. This contrasts with confocal activation, where optogenetic components above and below the focal plane (but inside the focal light cone) are activated as the system is typically highly light sensitive. Light-sheet illumination does result in some light scattering, but this effect is smaller than the light cone effect in confocal imaging. To reduce these effects, two-photon approaches have been developed ^9^. In addition, new, stable fluorescent proteins allow imaging of various cellular processes at the same time ^10^.

The Wnt pathway is a highly conserved signal transduction pathway functioning in development and disease in many organisms ^11^. In *Drosophila*, Wnt1 (Wingless or Wg) is a segment polarity gene that defines anterior-posterior patterning of the epidermis ^12^. Canonical Wnt signaling leads to inhibition of β-catenin (Armadillo or Arm in *Drosophila*) degradation, which subsequently activates downstream Wnt target genes in the nucleus. As β-catenin protein levels increase, it enters the nucleus where it regulates transcription of target genes by associating with the transcription factor TCF ^13,14^. This basic signaling mechanism is widely conserved, and critical in embryogenesis, but whether β-catenin signaling is needed in a constant widespread manner, versus in a temporally and/or spatially restricted manner was not known, because the tools needed to address this question were lacking. Here we describe our method for applying CRY2 protein fusions to investigate the spatial and temporal regulation of β-catenin protein in the developing embryo.

## Results

### Light-induced oligomerisation of β–catenin-CRY2-mCh modulates Wnt signaling in vitro

Blue light can induce oligomerization of *A. thaliana* CRY2 to form ‘photobodies’ in plant and mammalian cells ^6,15^. This leads to sequestration and hence functional inactivation of the protein. We tested this mechanism by fusing the photolyase homology region (PHR) of CRY2 and mCherry to the C terminus of *Drosophila* β–catenin or Arm protein (Arm-CRY2-mCh). To determine if the Arm-CRY2-mCh construct was responsive to light, we transfected it into S2R+ Wnt responsive *Drosophila* cells. Distinct fluorescent puncta were observed with blue light illumination (Fig. 1A; full construct details in Supplementary Fig. 1). The puncta dispersed, but did not entirely disappear upon withdrawal of blue light (Fig. 1A, B).

**Figure 1.**
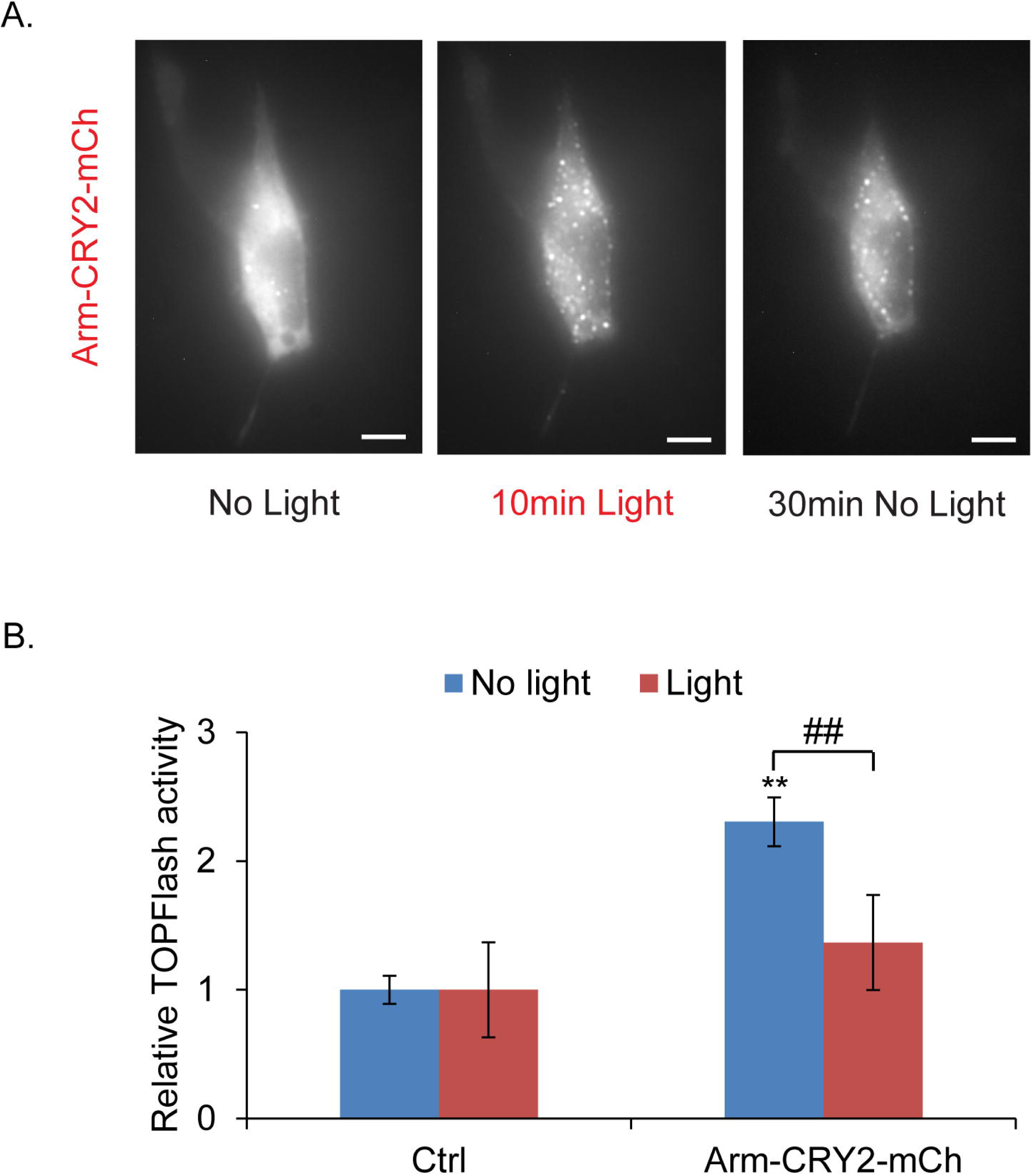
Light-induced clustering of Arm-CRY2-mCh modulates the Wnt/β-catenin pathway in *Drosophila*. (A) Arm-CRY2-mCh puncta formation in S2R+ cells in response to 488nm laser light. Scale bar represents 5µm. (B) Quantification of puncta in Arm-CRY2-mCh expressing S2R+ cells after illumination with 488nm laser light. One-way repeated measures ANOVA showed statistically significant difference in number of puncta between the different time points, ^##^p<0.01. (C) S2R+ cells transfected with pAct-Arm-CRY2-mCh plasmid and the TOPFlash luciferase reporter showed reduced TOPFlash activity after 16hr exposure to light. Graph is representative of three independent experiments and the average of three replicates (mean ± SD). The variance was found to be equal as determined by the F-test (p>0.05). Statistical significance relative to control samples was determined using the Student’s t-test. **p<0.01 relative to control, ^##^p<0.01. (D) Arm-CRY2-mCh puncta formation in *Drosophila* embryos overexpressing Arm-CRY2-mCh in a wild-type background upon exposure to 488nm laser light at 2.5min intervals. Puncta formation observed during illumination was reversible and oligomerization could be induced repeatedly. Scale bar represents 20µm. (E) Quantification of mean intensity (used as an indicator of puncta formation) in *Drosophila* embryos overexpressing Arm-CRY2-mCh in a wild-type background upon exposure to 488nm laser light at 2.5min intervals. The minimum occurring intensity was 210 as determined by the ZEN 2014 software and 225 was used as the lower threshold for detection of puncta formation. One-way repeated measures ANOVA showed statistically significant difference in mean intensity between the different time points,##p<0.01.

We next investigated if inducible Arm-CRY2-mCh puncta formation functionally inactivated Arm activity. The effect of the Arm-CRY2-mCh construct on canonical Wnt signaling was determined by using a TOPFlash reporter assay, where the luciferase gene is attached downstream of multimerized TCF binding sites ^13^. Overexpression of Arm-CRY2-mCh in S2R+ cells in the absence of light significantly increased TOPflash activity relative to control transfected cultures (Fig. 1C). This activation was inhibited when transfected cells were illuminated with light. Hence, oligomerization of the Arm-CRY2-mCh construct upon illumination with light effectively blocked downstream Wnt signaling.

Having verified the responsiveness of the Arm-CRY2-mCh construct to blue light illumination and its ability to modulate Wnt signaling *in vitro*, we tested the oligomerization of this construct in *Drosophila* embryos. Expression in wild type embryos showed the cytoplasmic and diffuse localization of Arm as indicated by the mCh fluorescence (Fig. 1D, Supplementary video 1). Since CRY2 has a turn-off speed of 5 minutes ^16^, the embryo was illuminated with blue light at 2.5 minute intervals at the lowest effective laser power (6%) and exposure time (7.5ms), to ensure continuous clustering while at the same time preventing excessive heating of the embryo. Upon blue light illumination, distinct fluorescent puncta were observed (Fig. 1D, Supplementary video 1) which plateaued 15 minutes after exposure (Fig. 1E). These results show an *in vivo*, light-responsive aggregation of β–catenin during development of the *Drosophila* embryo.

### *Arm-*CRY2*-mCh can rescue patterning in arm^XM19^ mutant embryos*

In order to establish the ability of the Arm-CRY2-mCh construct to function in place of endogenous Arm, we expressed it in an *arm*^*XM19*^ mutant background. We used *arm*^*XM19*^ mutants, because these lack the C terminus required for transcriptional transactivation but do not lose adhesion (Fig. 2A) ^17-19^. The basic test was to replace the mutated *arm*^*XM19*^ with Arm-CRY2-mCh (*arm*^*XM19*^/FM6; armGal4 crossed to UAS-Arm-CRY2-mCh). As the *arm* gene is located on the X-chromosome, normal, non-balancer, males from this cross can only be the rescued progeny. We observed normal males eclosing (three separate experiments, 31 of 64 males) from vials kept entirely in the dark and no normal males appearing from vials left out in normal room light. This genetic rescue experiment showed that Arm-CRY2-mCh could rescue *arm* mutants to viability as long as the flies were not exposed to light.

**Figure 2.**
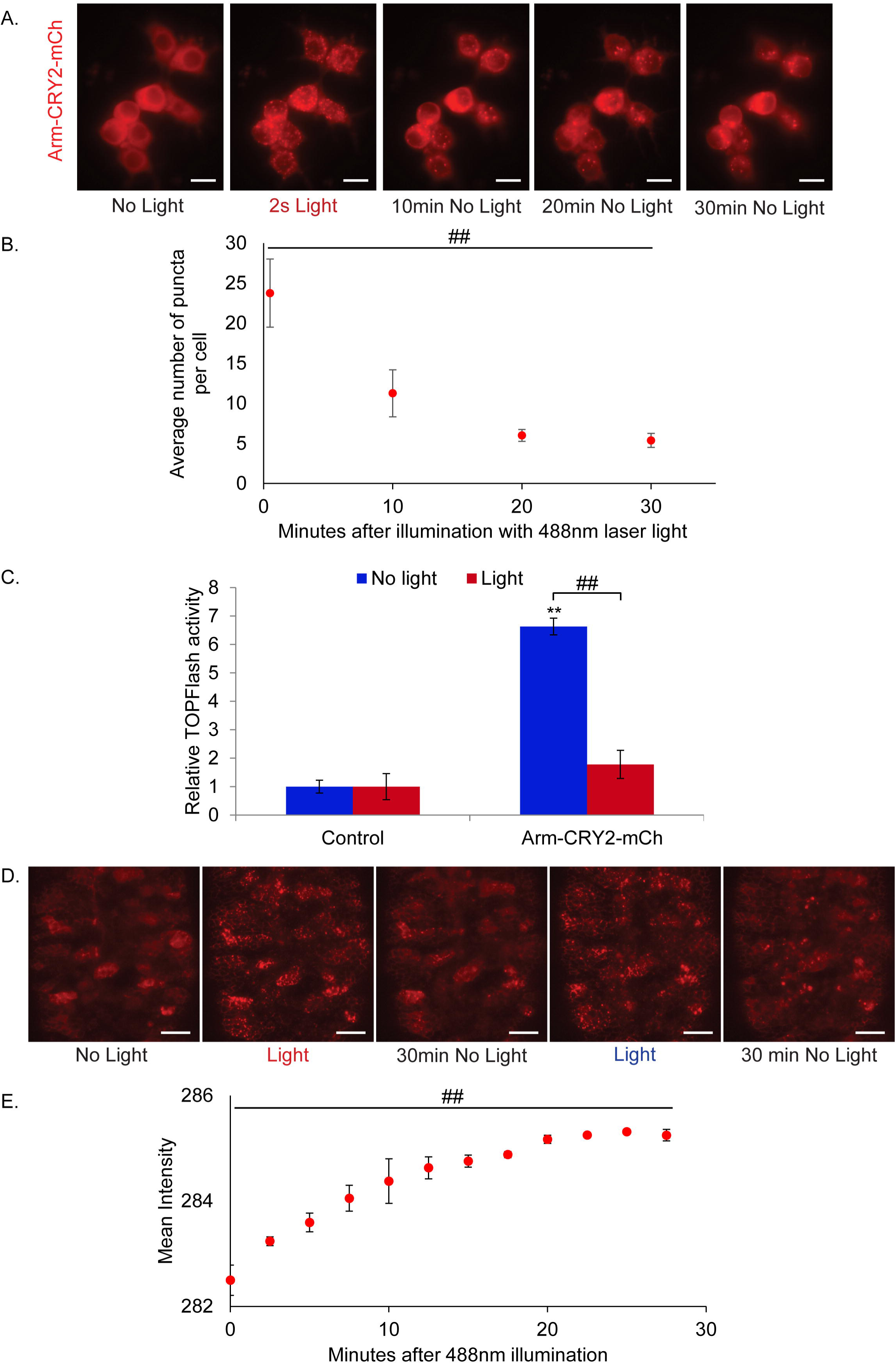
Arm-CRY2-mCh rescue of *arm*^*XM19*^ mutants is reversible with blue light. (A) Structure of *arm*^*XM19*^ truncated protein (adapted from ^48^. The N-terminus of Arm is required for transactivation, phosphorylation-based and proteasome-mediated degradation, and a-catenin binding. The central repeats contain the binding sites for most of Arm’s binding partners, such as APC, TCF, Cadherin, and Axin. The C-terminus is required for Cby and Teashirt binding and transactivation. *arm*^*XM19*^ lacks the entire C-terminus. (B) Cuticle preparations of *arm*^*XM19*^ mutant (germline clone, GLC) embryo with entire cuticle covered with denticles, *arm*^*XM19*^ GLC expressing Arm-CRY2-mCh without light was rescued to a wild-type cuticle (male embryos were identified by the presence of the *yellow* mutation and sorted away from the female embryos which were *y*^*+*^). *arm*^*XM19*^ GLC expressing Arm-CRY2-mCh with illumination showed defective Wg signaling (Denticle covered cuticle). Scale bar represents 50µm. (C) Quantitative expression of genes downstream of Wg signaling in Arm-CRY2-mCh overexpressing *arm*^*XM19*^ mutants. Pooled embryos exposed to light showed reduced *En* and *Wg* gene expression. Graph is representative of three independent experiments and the average of three replicates (mean ± SD). The variance was found to be equal as determined by the F-test (p>0.05). Statistical significance relative to control samples was determined using the Student’s t-test. *p<0.05, **p<0.01. (D) En antibody staining in Arm-CRY2-mCh overexpressing *arm*^*XM19*^ mutant embryos illuminated or not illuminated with light. En stripes as in wild-type embryos were observed in embryos not illuminated with light whereas embryos illuminated with light showed only small patches of En expression. Scale bar represents 50µm. (E) Stills from live imaging movies of Arm-CRY2-mCh overexpressing *arm*^*XM19*^ mutants. Embryos not subjected to light showed diffused localization of Arm-CRY2-mCh (inset) as compared to embryos subjected to light (inset). Embryos not subjected to light showed normal development whereas embryos subjected to light showed various developmental defects. Scale bar represents 50µm.

When canonical Wnt signaling is turned off, all ventral epidermal cells produce denticles. The opposite is true when Wnt signaling is turned on ectopically; most cells do not make denticles causing the naked phenotype (For a review of embryonic patterning ^20^). We made germline clones of *arm*^*XM19*^ where the maternal and zygotic contribution of protein was removed ^21^, resulting in an embryonic phenotype identical to *wg* mutant embryos (Fig. 2B)12,19,22. In these embryos, we expressed Arm-CRY2-mCh and saw a rescue of the canonical signaling defect (Fig. 2B) where *arm*^*XM19*^ mutants with Arm-CRY2-mCh showed a nearly wild type denticle pattern as long as the embryos were kept in the dark (Fig. 2B). When exposed to light, we observed no rescue of the canonical signaling defect as embryos showed the null signaling phenotype (Fig. 2B). Taken together, these results suggest that the Arm-CRY2-mCh construct can effectively rescue canonical Wnt signaling in *arm*^*XM19*^ mutant embryos, and that this rescue can be blocked by exposure to light.

To confirm the Wnt dependence of the phenotype, we examined genes downstream of Wnt signaling using quantitative PCR. Expression of the embryonic Wnt target genes *en* and *wg* was significantly decreased in Arm-CRY2-mCh expressing *arm*^*XM19*^ mutants exposed to light as contrasted with embryos kept in the dark both quantitatively (Fig. 2C), and qualitatively through immunostaining for En protein (Fig. 2D). In the absence of light, stripes of En protein were observed but these were disrupted with blue light illumination (Fig. 2D).

Next, we used light-sheet microscopy to live-image these embryos. *arm*^*XM19*^ mutants expressing the Arm-CRY2-mCh construct (as observed by luminescence upon excitation with the 561nm laser), showed normal embryonic development (Fig. 2E, Supplementary video 2A) similar to embryos expressing Arm-GFP (Supplementary video 2C). To ensure that the embryos imaged were the correct mutant (*arm*^*XM19*^ male embryos expressing the Arm-CRY2-mCh) we cuticle prepped the embryos post-imaging and scored for the *yellow* mutation. Only *y*^*-*^ embryos expressing mCh are presented here. We next exposed these embryos to blue light and observed smaller embryos developing with a lawn of denticles (Fig. 2E, Supplementary video 2B), consistent with a loss of Wnt signaling. A troubling finding was that the puncta once formed did not disappear quickly raising the possibility that the effect was not reversible. To test this, we performed experiments where embryos were placed in the light sheet microscope and illuminated with blue light every 2.5, 5 or 10 minutes. *arm*^*XM19*^ mutants expressing the Arm-CRY2-mCh construct illuminated every 2.5 or 5 minutes did not survive to the end of embryogenesis, but those illuminated only once every 10 minutes hatched. We propose that the persistence of puncta represents the cells’ inability to degrade aggregated Arm protein quickly, but this does not affect Wnt signaling as enough Arm protein is made during the dark phase to compensate for the aggregates as long as 10 minutes are allowed to elapse. The system, therefore, is reversible but may require new protein synthesis rather than dis-aggregation.

### Temporal regulation of Wnt signaling during development in arm^XM19^ embryos

Having established the efficacy of the Arm-CRY2-mCh transgene in embryos, we next performed a temporal experiment to answer the biological question of when Wnt signaling is required. Segment polarity genes are expressed downstream of pair rule genes early in development ^22^. These genes turn on Wg expression ^23^ through a regulatory loop with Engrailed ^24,25^. Experiments with a temperature sensitive allele of *wg* established four phases of Wg signaling in establishment and maintenance of embryonic patterning ^26^. These can be subdivided into early *en* activation and later *shaven baby* repression in combination with EGF signals ^24,27,28^. We used eggs exposed to light at different developmental stages ^29^, and observed the cuticle phenotypes. Staging was approximate as the experiment was carried out at 21°C. Exposure to light from stages 1 to 11 (From cleavage stages to Germ band elongation) showed a variable lawn of denticles similar to *wg* mutants (Fig. 3A). Strong Wnt pathway loss of function embryos show a lawn of denticles and are much smaller than wild-type. Restoring some Wnt pathway function rescues the small size phenotype ^12,30^. The lawn of denticles began getting broader (size of the embryo increased) when light was applied at stage 12 (onset germ band retraction) with some distance developing between the denticle bands (Fig. 3A, stage 13 germ band retraction). This experiment showed that Arm-CRY2-mCh could be used to recapitulate previous findings with *wg*^*ts*^ that showed effects into dorsal closure stages ^26^.

**Figure 3.**
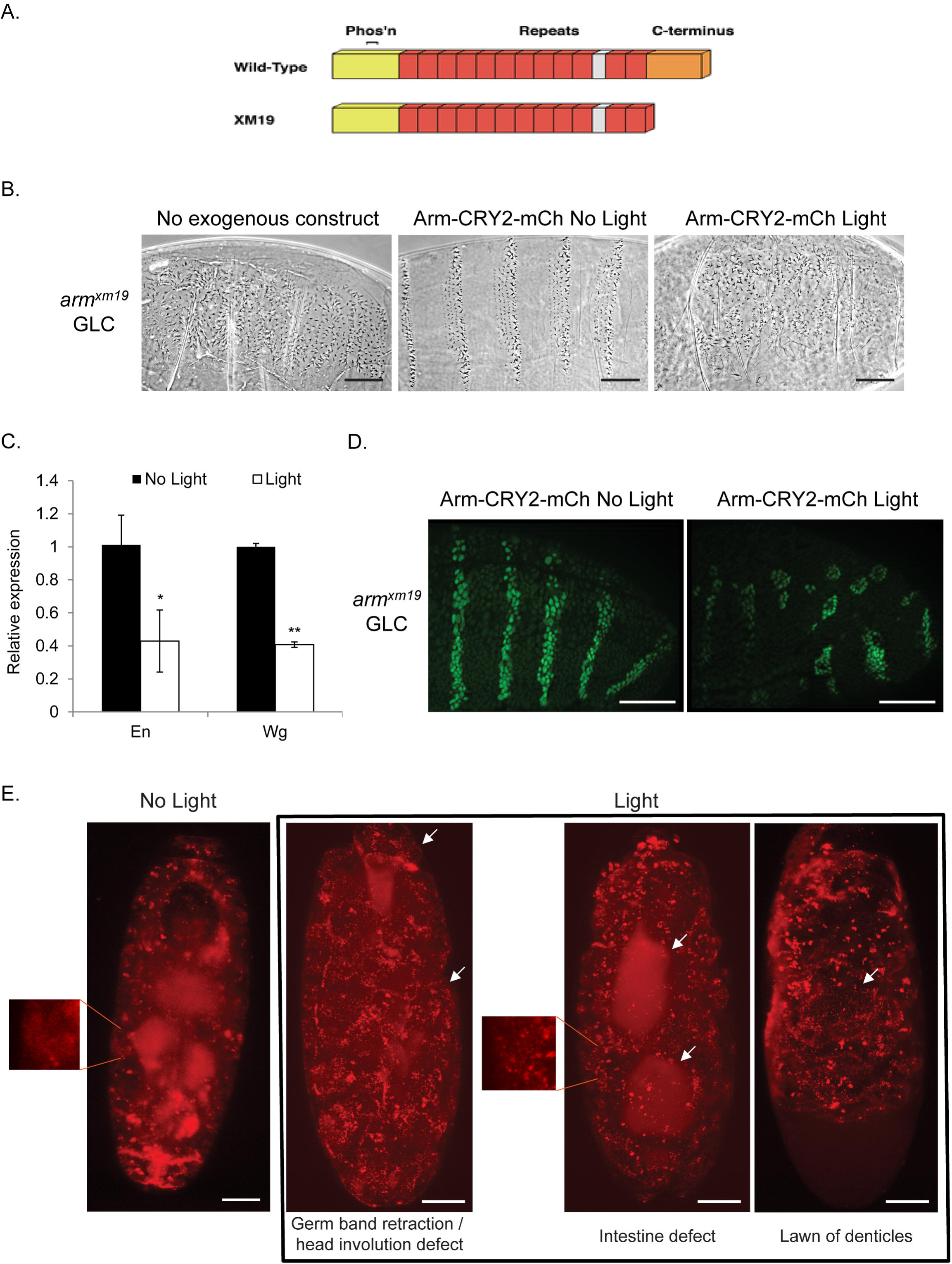
Temporal and spatial inactivation of Arm-CRY2-mCh in *arm*^*XM19*^ mutant embryos overexpressing Arm-CRY2-mCh. (A) *arm*^*XM19*^ mutant embryos overexpressing Arm-CRY2-mCh were collected and exposed to light at various developmental stages. Cuticle preparations were carried out after light exposure. Exposure to light earlier (stage 1 to 11) during development resulted in a *wg* mutant phenotype. The denticle patterning began spreading out thereafter until stage 13 after which a mostly wild-type phenotype was observed. Scale bar represents 50µm. (B) Spatial perturbation of Wnt signaling in Arm-CRY2-mCh overexpressing *arm*^*XM19*^ mutant embryos along line of symmetry. The right half of the embryo was illuminated with 488nm laser light and the whole embryo was imaged simultaneously using light sheet microscopy. The right side of the embryo (with blue light illumination) showed distinct clustering and developmental defects. Scale bar represents 50µm. (C) Spatial perturbation of Wnt signaling in Arm-CRY2-mCh overexpressing *arm*^*XM19*^ mutant embryos along anterior-posterior axis. The posterior region of the embryo was illuminated with 488nm laser light and the whole embryo was imaged simultaneously using light sheet microscopy. The posterior portion of the embryo (with blue light illumination) showed distinct clustering and incomplete germ band retraction. Scale bar represents 50µm.

### Spatial perturbation of Wnt signaling during development in arm^XM19^ embryos

We next wanted to test the spatial resolution of this approach using light sheet microscopy as a blue light source as well as a recorder for live imaging. Light sheet microscopy is not an ideal tool for this experiment as the light is not delivered as a point (For a better approach to spatial resolution see ^31^). Half of an embryo overexpressing the Arm-CRY2-mCh transgene was illuminated with blue light and the whole embryo imaged in the red channel. The region of the embryo with blue light illumination showed similar defects as in the earlier results with the unilluminated half remaining more wild-type (Fig. 3B, Supplementary Video 3A, compare to Supplementary Video 2A and 2B). Spatial manipulation of signaling was further tested by illuminating a portion of the embryo along the anterior-posterior axis. Illumination of the posterior region of the embryo showed distinct clustering and developmental defects as compared to the anterior portion of the embryo (Fig. 3C, Supplementary Video 3B). These results show that combining light sheet imaging with spatial inactivation of Wnt signaling can be used to make and image mosaic embryos.

### *Light-responsive properties of Arm-*CRY2*-mCh can modulate Wnt signaling in HEK293*

We have shown that Arm-CRY2-mCh functions in insect cells and in *Drosophila* embryos. We next tested if this construct could be applied to other experimental systems by looking at whether it could be used to modulate Wnt signaling in mammalian cells. We used HEK293 (Human embryonic kidney cells) stably transfected with the Wnt3a gene and TOPFlash vector (HEK293-Wnt3a-TOPFlash). Cells transfected with the construct showed fluorescent puncta formation upon blue light illumination (Fig. 4A). The puncta dispersed rapidly upon withdrawal of blue light (Fig. 4A).

**Figure 4.**
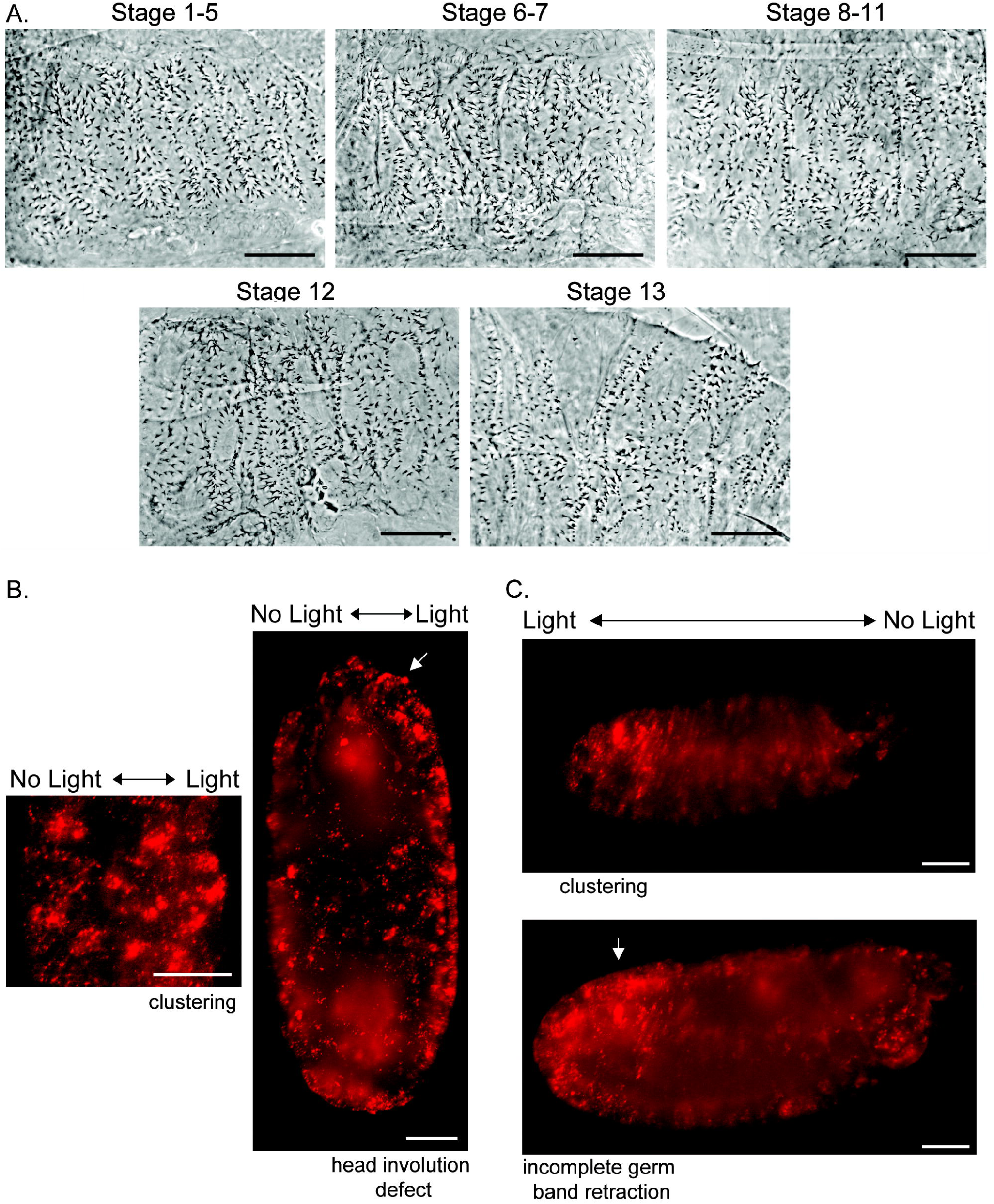
Light-induced clustering of Arm-CRY2-mCh modulates the Wnt/β-catenin pathway in mammalian cells. (A) Arm-CRY2-mCh puncta formation in HEK293-Wnt3a-TOPFlash cells in response to 488nm laser light, as well as dissociation after light withdrawal. Scale bar represents 10µm. (B) HEK 293-Wnt3a-TOPFlash cells transfected with pAct-Arm-CRY2-mCh plasmid and the TOPFlash luciferase reporter showed reduced TOPFlash activity after exposure to light. Graph is representative of three independent experiments and the average of three replicates (mean ± SD). The variance was found to be equal as determined by the F-test (p>0.05). Statistical significance relative to control samples was determined using the Student’s t-test. **p<0.01 relative to control, ^##^p<0.01.

The effect of inducible Arm-CRY2-mCh clustering on Wnt signaling in mammalian cells was quantified using a TOPFlash reporter assay. Overexpression of Arm-CRY2-mCh plasmid in HEK293-Wnt3a-TOPFlash cells in the absence of light significantly increased TOPflash activity relative to control transfected cultures (Fig. 4B). This activation was inhibited when transfected cells were exposed to light. Our data suggest that oligomerization of the Arm-CRY2-mCh construct upon illumination with light could be applied to other experimental systems.

## Discussion

Classical genetics tools allow tissue-specific manipulation of gene expression, but not precise temporal or spatial manipulation. Optogenetic tools can address this problem by allowing spatial and temporal regulation in a rapid, precise, and reversible manner ^2,32,33^. Various optogenetic systems have been described to date such as the Cryptochrome 2 (CRY2), Phytochrome B (PHYB), LOV and Dronpa systems. However, only the CRY2 and LOV systems allow optogenetic control at a lower wavelength with live imaging at a higher wavelength. Both systems are activated within a few seconds, however, the CRY2 system has a slower turn off speed of approximately 5 minutes as compared to tens of seconds or minutes in the case of LOV ^16^. This makes the CRY2 system ideal for live imaging with lower phototoxicity. Our study is the first to combine optogenetics with light-sheet microscopy for noninvasive, temporal and spatial control of cell signaling *in vivo*. Using these tools, we investigated the spatial and temporal regulation of β–catenin and Wnt signaling *in vivo* and *in vitro* proving the efficacy of this method. Wnt’s role in segment polarity is well known ^34,35^, and we observed the same phenotypes by inactivating Wnt signaling during early stages of development (stage 1 to 11). Light-sheet microscopy is not designed for single cell conversion of optogenetic switches ^31^, but can be applied to sections of the embryo to study problems such as left-right asymmetry and anterior-posterior patterning by illuminating different regions of the embryo. Other microscopy approaches can be used to prevent light scattering ^9^, where the optical molecule could be activated in unintended parts of the embryo, but the advantage of light sheet microscopy is that it allows whole embryo imaging over the entire course of the 24 hour Drosophila embryogenesis.

Wnt signaling in segment polarity is well studied, but many additional functions have been observed ^36^. Recently, ^37^ reported that intestinal formation during embryo development was disrupted in embryos treated with ionomycin, an inhibitor of Wnt/β-catenin signaling. By coupling optogenetics with live imaging, we observed a similar defect in intestinal development in Arm-CRY2-mCh expressing *arm*^*XM19*^ mutant embryos. Our experiment with spatial inactivation of Arm-CRY2-mCh in only a specified region of the embryo, also confirmed these findings. We see the application of optogenetics with light-sheet microscopy will allow the exploration of many more Wnt pathway effects. The main limitation appears to be the buildup of CRY2 puncta that do not disaggregate. In our experiments this leads to a buildup of mCh fluorescence in the later stages especially in hemocyte cells making imaging in the red channel difficult at late stages. We anticipate that this can be overcome by using separate fluorescent proteins for imaging and for optogenetics.

Our optogenetics approach is a non-invasive and versatile tool that allows for manipulation as well as live tracking of protein activity to interrogate complex phenotypes *in vivo* ^38^. We foresee applying this tool to cell culture, organoid studies, vertebrate development in zebrafish, and many other systems ^39^. It provides an alternative to small molecule studies which can only be performed on druggable targets. We have made a human β–catenin-CRY2 version that functions similarly, and are applying the optogenetic methods to study adhesion by using embryos with a stronger loss of function *arm*^*O43A01*^ allele that disrupts junctions but is rescued by Arm-CRY2-mCh ^40-42^. Further, we find that NFκB/Dorsal and Erk pathways are also effectively studied in this way as our Dorsal-CRY2 and Ras-CRY2 constructs function well in tissue culture and hopefully soon in flies.

## Materials and Methods

### Transgenes and GAL4 driver lines

Two ubiquitous drivers were used for expression of transgenes: the weaker armadillo-GAL4 and the stronger daughterless-GAL4 ^43^. The Arm-CRY2-mCh constructs were made using the MultiSite Gateway^®^ Pro 2.0 cloning kit (Invitrogen). Full-length Armadillo was amplified from *Drosophila* embryo cDNA with *att*B1 and *att*B5r-flanked sites. This PCR product was recombined into pDONR P1-P5r vector. The CRY2-mCh was amplified from pCRY2PHR-mCherryN1 which was a gift from Chandra Tucker (Addgene plasmid # 26866)6. *att*B5 and *att*B2 sites were added to the CRY2-mCh PCR product and recombined into pDONR P5-P2 vector. Both pDONR vectors with Armadillo and CRY2-mCh insert were recombined by Gateway cloning (Invitrogen) into pUASg.attB with COOH-terminal 3XHA tag (A kind gift from J. Bischof and K. Basler, Zurich)^44^ to obtain the Arm-CRY2-mCh transgene for expression in *Drosophila*. Transgenes were injected into attP2 (Strain #8622) P[CaryP]attP2 68A4 by BestGene Inc. (California)^45^. For constitutive expression in S2R+ cells, the pDONR vectors were recombined into the Gateway destination vector, pAW (Drosophila Gateway^®^ Vector collection, Carnegie Institution). For constitutive expression in mammalian cell lines, the pDONR vectors were recombined into the mammalian Gateway destination vector, Gateway pcDNA-pDEST40 (ThermoFisher Scientific, USA). EGFP-tagged protein lines were derived using the *Minos* mediated integration cassette (MiMIC) methodology described by Nagarkar-Jaiswal et al. ^46,47^ and the Arm MiMIC line was obtained from the Bloomington Drosophila Stock Center (BDSC).

### Crosses and expression of UAS constructs

The dominant female sterile technique was used to generate maternally mutant eggs ^21^. Please see Flybase for details on mutants used (flybase.bio.indiana.edu). For mis-expression experiments, the ArmGAL4 2^nd^ chromosome was used. FRT 101 was used to generate the X chromosome mutant.

The following cross was conducted to generate germline clone embryos: *y*^*-*^, *arm*^*XM19*^ FRT101/ovoD1 FRT101; Arm-Gal4 x *w*^*-*^, *y*^*+*^; UAS-Arm-CRY2-mCh Embryos were imaged in the light sheet microscope showing mCh expression, and were further cuticle prepped after imaging to confirm the genotype and phenotype as described for *y*^*-*^, arm^XM19^ mutation, an assay based on the pigmentation of the denticles ^40,41,48^.

The following cross was conducted to generate zygotic mutant rescue flies: *y*^*-*^, *arm*^*XM19*^ FRT101/FM6; Arm-Gal4 x *w*^*-*^, *y*^*+*^/Y-chromosome; UAS-Arm-CRY2-mCh In this cross, all males receive the X-chromosome from their mothers so only balancer (FM6, *Bar*) males are expected unless the transgene can rescue the lethality of *arm*^*XM19*^. Identical crosses were set up with three kept entirely in the dark and three under normal laboratory light conditions for two weeks. The number and phenotype of males from all the crosses was scored.

### Light-sheet microscopy

Embryos were dechorionated in bleach, washed in water and dried on a paper towel. They were then embedded into a capillary containing 1% agarose, low gelling temperature Type VII-A (Sigma) dissolved in water, using a fine wire such that the embryo was upright with the anterior-posterior axis aligned with the axis or perpendicular to the axis of the glass capillary. For imaging, the agarose was pushed out of the capillary and the sample was suspended freely in the sample chamber containing water as our lightsheet set-up utilized a water immersion objective.

Live imaging of *Drosophila* embryos was carried out on the light-sheet Z.1 fluorescence microscope (Carl Zeiss, Germany) with the Lightsheet Z.1 10x/0.2 Illumination Optics and W Plan-Apochromat 20×/1.0 UV-VIS detection objective (Carl Zeiss, Germany). The lightsheet microscope was equipped with the Lightsheet Z.1 detection module “Standard”, 30mW 488nm and 20mW 561nm solid state lasers with BP505-545 and LP585 emission filters respectively. Whole embryos (unless otherwise stated) were imaged using dual-side illumination by a light-sheet modulated into a pivot scan mode. The 488nm laser was used at 6% power with 7.5ms exposure time and the 561nm laser was used at 13% laser power with 12.5ms exposure time. Since the oligomerization is reversible with removal of blue light, the whole embryo was excited with both laser lines every 2.5 min in Z-stack mode at 1µm step size for 24 hrs. For control embryos not exposed to blue light, the same settings were used but with illumination using only the 561nm laser line. All single plane illumination (SPIM) data was saved in the LSM format and processed using the ZEN 2014 SP1 software (Carl Zeiss, Germany). Background fluorescence was automatically calculated by the ZEN 2014 software using the background correction function (ZEN Software Guide, Carl Zeiss, Germany). Z-stacks that were deep in the embryo and out of focus were removed and maximum intensity projections of z-stacks in focus were generated for each embryo ^49^. Images for figures were processed and assembled using Photoshop and Illustrator programs from Adobe.

### Antibodies and Immunofluorescence

Embryos were dechorionated in bleach and fixed with heptane/4% formaldehyde in phosphate buffer (0.1M NaPO4 pH 7.4) for 20 minutes ^40^. The aqueous phase was removed and an equal amount of methanol was added to devitellinize the embryos. Antibody stainings were done in PBT (PBS, 0.1% Triton X-100, 1% bovine serum albumin, 0.1% Azide). The anti-Engrailed antibody mAbs (4D9) was obtained from the Developmental Studies Hybridoma Bank, created by the NICHD of the NIH and maintained at The University of Iowa, Department of Biology, Iowa City, IA 52242 and used at a dilution of 1:10. Alexa Fluor 488-anti-mouse secondary antibody was used (ThermoFisher Scientific, USA) at a 1:1000 dilution. Embryos were mounted in Aquapolymount^®^ (Polysciences, Inc.). Images were acquired on the Zeiss AxioImager Z1 with Apotome (Carl Zeiss, Germany) and images were processed using the AxioVision Rel. 4.8 (Carl Zeiss, Germany).

### Live-cell imaging.

Time-lapse microscopy of activated CRY2 fusions in S2R+ and endogenously expressing WNT3A in a HEK 293 cell line that contains an integrated TOPFlash reporter (A kind gift from Dr Alan Prem Kumar, Cancer Science Institute of Singapore) was performed using a Zeiss AxioImager Z1 microscope (Carl Zeiss, Germany). Clustering visualization was carried out at room temperature. Blue light exposure and mCherry imaging were performed simultaneously by imaging in both 488nm and 561nm laser channels. Images were acquired and processed using the AxioVision Rel. 4.8 (Carl Zeiss, Germany). Puncta counts were determined using the Cell Counter plugin in ImageJ.

### TOPflash assay

TOPflash luciferase assays (TCF/LEF reporter assays) were performed to assess the effect of light on canonical Wnt-signaling in cells transfected with the Arm-CRY2-mCh plasmid. S2R+ cells were co-transfected with TOPflash ^13^, *Renilla* luciferase-Pol III (Renilla luciferase-Pol III was a gift from Norbert Perrimon (Addgene plasmid # 37380)) ^50^ and the pAW-Arm-CRY2-mCh plasmid using lipofectamine 3000 (ThermoFisher Scientific) according to the manufacturer’s instructions. In the case of the HEK 293 cell line endogenously expressing WNT3A and the TOPFlash reporter, *Renilla* luciferase expressing vector (pRL-CMV, Promega, USA) and the pDEST40-Arm-CRY2-mCh plasmid were transfected using lipofectamine 3000 (ThermoFisher Scientific) according to the manufacturer’s instructions. Cell lysates were prepared 24 hours after transfection and luciferase activity was measured using the Dual-Luciferase Reporter Assay System (Promega) according to the manufacturer’s instructions. The relative TOPflash luciferase activity was measured using the ratio of firefly/renilla luciferase activity and the data was presented as mean ± SD. Statistical analyses were performed using Student’s *t* test unless the data showed evidence of unequal standard deviation (F-test, *p* < 0.05). In these cases, the Mann–Whitney nonparametric test was used.

### RNA Extraction, cDNA Synthesis and qPCR

Mixed stage and genotype embryos, 0 to 16 hours after deposition, were exposed to light or no light for 6 hours, dechorionated in bleach, washed in water and collected. Total RNA was extracted for each treatment using the RNeasy Mini Kit (Qiagen) as per the manufacturer’s protocol. Total RNA concentration and purity was measured using the Cytation 3 Cell Imaging Multi-Mode Reader (BioTek). One µg of total RNA was reverse transcribed in a 20µl reaction volume using the QuantiTect reverse transcription kit (Qiagen) according to the manufacturer’s protocol. Gene specific primer sequences were obtained from Fly Primer Bank ^51^. Quantification of mRNA was performed using SYBR^®^ Green Assay (Thermo Fisher Scientific) on the PikoReal™ Real-Time PCR System (Thermo Fisher Scientific) and a PCR product dissociation curve was generated to ensure specificity of amplification. For expression analysis, the mRNA data were normalized to the endogenous control, RPL32, followed by calibration to cultures not exposed to light using relative quantification (2^-^ΔΔCT).Results were generated from 3 technical replicates for each mRNA. The average relative expression ± standard deviation (SD) was determined. Statistical analyses were performed using Student’s *t* test unless the data showed evidence of unequal standard deviation (F-test, *p* < 0.05). In these cases, the Mann–Whitney nonparametric test was used.

### Contributions

P.K. and N.S.T. designed and performed the experiments. N.S.T., P.K. and T.E.M. wrote the paper. All authors reviewed manuscript.

## Acknowledgments

We thank Yale-NUS students, Le Van Canh, Hyung-Seok Kim and Xiao Linfan for their help. This work was supported by an Academic Research Fund (AcRF) grant (MOE2014-T2-2-039) of the Ministry of Education, Singapore to NST. TES was supported by a National Research Foundation NRF Fellowship (NRF2012NRF-NRFF001-094).

## Supplementary Videos Legends

**Supplementary Video 1.** Arm-CRY2-mCh puncta formation in *Drosophila* embryos overexpressing Arm-CRY2-mCh in a wild-type background upon exposure to 488nm laser light. The whole embryo was illuminated with 488nm laser light at 2.5 minute intervals for half an hour to activate oligomerization. Puncta formation observed during illumination was reversible and oligomerization could be induced repeatedly.

**Supplementary Video 2A.** Arm-CRY2-mCh overexpressing *arm*^*XM19*^ mutant embryos not exposed to light showed diffuse localization of Arm-CRY2-mCh and normal embryonic development.

**Supplementary Video 2B.** Arm-CRY2-mCh overexpressing *arm*^*XM19*^ mutant embryos exposed to 488nm light showed distinct puncta formation of Arm-CRY2-mCh and various developmental defects as compared to embryos not exposed to light.

**Supplementary Video 2C.** Embryos expressing Arm-EGFP showed normal embryonic development.

**Supplementary Video 3A.** Spatial perturbation of Wnt signaling in Arm-CRY2-mCh overexpressing *arm*^*XM19*^ mutant embryos. The right half of the embryo was illuminated with 488nm laser light and the whole embryo was imaged simultaneously in the red channel. The right side of the embryo (with blue light illumination) showed distinct clustering and developmental defects.

**Supplementary Video 3B.** Spatial perturbation of Wnt signaling in Arm-CRY2-mCh overexpressing *arm*^*XM19*^ mutant embryos. The posterior region of the embryo was illuminated with 488nm laser light and the whole embryo was imaged simultaneously in the red channel. The posterior region of the embryo (with blue light illumination) showed distinct clustering and incomplete germ band retraction.

## References

1 Doupe, D. P. & Perrimon, N. Visualizing and manipulating temporal signaling dynamics with fluorescence-based tools. Sci Signal 7, re1, doi:10.1126/scisignal.2005077 (2014).

2 Aravanis, A. M. et al. An optical neural interface: in vivo control of rodent motor cortex with integrated fiberoptic and optogenetic technology. J Neural Eng 4, S143–156, doi:10.1088/1741-2560/4/3/S02 (2007).

3 Bugaj, L. J., Choksi, A. T., Mesuda, C. K., Kane, R. S. & Schaffer, D. V. Optogenetic protein clustering and signaling activation in mammalian cells. Nat Methods 10, 249–252, doi:10.1038/nmeth.2360 (2013).

4 Liu, H., Gomez, G., Lin, S., Lin, S. & Lin, C. Optogenetic control of transcription in zebrafish. PLoS One 7, e50738, doi:10.1371/journal.pone.0050738 (2012).

5 Konermann, S. et al. Optical control of mammalian endogenous transcription and epigenetic states. Nature 500, 472–476, doi:10.1038/nature12466 (2013).

6 Kennedy, M. J. et al. Rapid blue-light-mediated induction of protein interactions in living cells. Nat Methods 7, 973–975, doi:10.1038/nmeth.1524 (2010).

7 Huang, A., Amourda, C., Zhang, S., Tolwinski, N. S. & Saunders, T. E. Decoding temporal interpretation of the morphogen Bicoid in the early Drosophila embryo. Elife 6, doi:10.7554/eLife.26258 (2017).

8 Huisken, J. & Stainier, D. Y. Selective plane illumination microscopy techniques in developmental biology. Development 136, 1963–1975, doi:10.1242/dev.022426 (2009).

9 Prakash, R. et al. Two-photon optogenetic toolbox for fast inhibition, excitation and bistable modulation. Nat Methods 9, 1171–1179, doi:10.1038/nmeth.2215 (2012).

10 Chudakov, D. M., Matz, M. V., Lukyanov, S. & Lukyanov, K. A. Fluorescent proteins and their applications in imaging living cells and tissues. Physiol Rev 90, 1103–1163,doi:10.1152/physrev.00038.2009 (2010).

11 Clevers, H. Wnt/beta-catenin signaling in development and disease. Cell 127, 469–480, doi:10.1016/j.cell.2006.10.018 (2006).

12 Rijsewijk, F. et al. The Drosophila homolog of the mouse mammary oncogene int-1 is identical to the segment polarity gene wingless. Cell 50, 649–657 (1987).

13 van de Wetering, M. et al. Armadillo coactivates transcription driven by the product of the Drosophila segment polarity gene dTCF. Cell 88, 789–799 (1997).

14 Brunner, E., Peter, O., Schweizer, L. & Basler, K. pangolin encodes a Lef-1 homologue that acts downstream of Armadillo to transduce the Wingless signal in Drosophila. Nature 385, 829–833 (1997).

15 Mas, P., Devlin, P. F., Panda, S. & Kay, S. A. Functional interaction of phytochrome B and cryptochrome 2. Nature 408, 207–211, doi:10.1038/35041583 (2000).

16 Tischer, D. & Weiner, O. D. Illuminating cell signalling with optogenetic tools. Nat Rev Mol Cell Biol 15, 551–558, doi:10.1038/nrm3837 (2014).

17 Wieschaus, E., Nusslein-Volhard, C. & Jurgens, G. Mutations affecting the pattern of the larval cuticle in Drosophila melanogaster. Roux Arch. dev. Biol. 193, 296–307 (1984).

18 Riggleman, B., Wieschaus, E. & Schedl, P. Molecular analysis of the armadillo locus: uniformly distributed transcripts and a protein with novel internal repeats are associated with a Drosophila segment polarity gene. Genes Dev 3, 96–113 (1989).

19 Peifer, M. & Wieschaus, E. The segment polarity gene armadillo encodes a functionally modular protein that is the Drosophila homolog of human plakoglobin. Cell 63, 1167–1176 (1990).

20 Hatini, V. & DiNardo, S. Divide and conquer: pattern formation in Drosophila embryonic epidermis. Trends Genet 17, 574–579 (2001).

21 Chou, T. B. & Perrimon, N. Use of a yeast site-specific recombinase to produce female germline chimeras in Drosophila. Genetics 131, 643–653 (1992).

22 Nusslein-Volhard, C. & Wieschaus, E. Mutations affecting segment number and polarity in Drosophila. Nature 287, 795–801 (1980).

23 Ingham, P. W., Baker, N. E. & Martinez-Arias, A. Regulation of segment polarity genes in the Drosophila blastoderm by fushi tarazu and even skipped. Nature 331, 73–75, doi:10.1038/331073a0 (1988).

24 Heemskerk, J., DiNardo, S., Kostriken, R. & O'Farrell, P. H. Multiple modes of engrailed regulation in the progression towards cell fate determination. Nature 352, 404–410, doi:10.1038/352404a0 (1991).

25 DiNardo, S., Sher, E., Heemskerk-Jongens, J., Kassis, J. A. & O'Farrell, P. H. Two-tiered regulation of spatially patterned engrailed gene expression during Drosophila embryogenesis. Nature 332, 604–609, doi:10.1038/332604a0 (1988).

26 Bejsovec, A. & Martinez Arias, A. Roles of wingless in patterning the larval epidermis of Drosophila. Development 113, 471–485 (1991).

27 Payre, F., Vincent, A. & Carreno, S. ovo/svb integrates Wingless and DER pathways to control epidermis differentiation. Nature 400, 271–275, doi:10.1038/22330 (1999).

28 Sanson, B., Alexandre, C., Fascetti, N. & Vincent, J. P. Engrailed and hedgehog make the range of Wingless asymmetric in Drosophila embryos. Cell 98, 207–216 (1999).

29 Hartenstein, V. Atlas of Drosophila development. (Cold Spring Harbor Laboratory Press, 1993).

30 Suresh, J. et al. An embryonic system to assess direct and indirect Wnt transcriptional targets. Scientific Reports 7, 11092, doi:10.1038/s41598-017-11519-z (2017).

31 Johnson, H. E. et al. The Spatiotemporal Limits of Developmental Erk Signaling. Dev Cell 40,185–192, doi:10.1016/j.devcel.2016.12.002 (2017).

32 Galvan, A., Hu, X., Smith, Y. & Wichmann, T. In vivo optogenetic control of striatal and thalamic neurons in non-human primates. PLoS One 7, e50808,doi:10.1371/journal.pone.0050808 (2012).

33 Jia, Z. et al. Stimulating cardiac muscle by light: cardiac optogenetics by cell delivery. Circ Arrhythm Electrophysiol 4, 753–760, doi:10.1161/CIRCEP.111.964247 (2011).

34 Willert, K. & Nusse, R. Beta-catenin: a key mediator of Wnt signaling. Curr Opin Genet Dev 8,95–102, doi:S0959-437X(98)80068-3 [pii] (1998).

35 Klingensmith, J. & Nusse, R. Signaling by wingless in Drosophila. Dev Biol 166, 396–414,doi:10.1006/dbio.1994.1325 (1994).

36 Swarup, S. & Verheyen, E. M. Wnt/Wingless signaling in Drosophila. Cold Spring Harb Perspect Biol 4, doi:10.1101/cshperspect.a007930 (2012).

37 Akiyoshi, R., Kaneuch, T., Aigaki, T. & Suzuki, H. Bioluminescence imaging to track real-time armadillo promoter activity in live Drosophila embryos. Anal Bioanal Chem 406, 5703–5713, doi:10.1007/s00216-014-8000-8 (2014).

38 Shin, Y. et al. Spatiotemporal Control of Intracellular Phase Transitions Using Light-Activated optoDroplets. Cell 168, 159–171 e114, doi:10.1016/j.cell.2016.11.054 (2017).

39 Krzic, U., Gunther, S., Saunders, T. E., Streichan, S. J. & Hufnagel, L. Multiview light-sheet microscope for rapid in toto imaging. Nat Methods 9, 730–733, doi:10.1038/nmeth.2064 (2012).

40 Tolwinski, N. S. & Wieschaus, E. Armadillo nuclear import is regulated by cytoplasmic anchor Axin and nuclear anchor dTCF/Pan. Development 128, 2107–2117 (2001).

41 Tolwinski, N. S. et al. Wg/Wnt signal can be transmitted through arrow/LRP5,6 and Axin independently of Zw3/Gsk3beta activity. Dev Cell 4, 407–418 (2003).

42 Kaplan, N. A., Liu, X. & Tolwinski, N. S. Epithelial polarity: interactions between junctions and apical-basal machinery. Genetics 183, 897–904, doi:10.1534/genetics.109.108878 (2009).

43 Brand, A. H. & Perrimon, N. Targeted gene expression as a means of altering cell fates and generating dominant phenotypes. Development 118, 401–415 (1993).

44 Bischof, J. et al. A versatile platform for creating a comprehensive UAS-ORFeome library in Drosophila. Development 140, 2434–2442, doi:10.1242/dev.088757 (2013).

45 Groth, A. C., Fish, M., Nusse, R. & Calos, M. P. Construction of transgenic Drosophila by using the site-specific integrase from phage phiC31. Genetics 166, 1775–1782 (2004).

46 Nagarkar-Jaiswal, S. et al. A genetic toolkit for tagging intronic MiMIC containing genes. Elife 4, doi:10.7554/eLife.08469 (2015).

47 Venken, K. J. et al. MiMIC: a highly versatile transposon insertion resource for engineering Drosophila melanogaster genes. Nat Methods 8, 737–743 (2011).

48 Tolwinski, N. S. & Wieschaus, E. A nuclear function for armadillo/beta-catenin. PLoS Biol 2, E95 (2004).

49 Novak, D., Kucharova, A., Ovecka, M., Komis, G. & Samaj, J. Developmental Nuclear Localization and Quantification of GFP-Tagged EB1c in Arabidopsis Root Using Light-Sheet Microscopy. Front Plant Sci 6, 1187, doi:10.3389/fpls.2015.01187 (2015).

50 Nybakken, K., Vokes, S. A., Lin, T. Y., McMahon, A. P. & Perrimon, N. A genome-wide RNA interference screen in Drosophila melanogaster cells for new components of the Hh signaling pathway. Nat Genet 37, 1323–1332, doi:10.1038/ng1682 (2005).

51 Hu, Y. et al. FlyPrimerBank: an online database for Drosophila melanogaster gene expression analysis and knockdown evaluation of RNAi reagents. G3 (Bethesda) 3, 1607–1616, doi:10.1534/g3.113.007021 (2013).

